# Proteomic profiling of regenerated urinary bladder tissue with stem cell seeded scaffold composites in a non-human primate bladder augmentation model

**DOI:** 10.1101/2023.08.29.554824

**Authors:** Tiffany T. Sharma, Seby L. Edassery, Natchiket Rajinikanth, Vikram Karra, Matthew I. Bury, Arun K. Sharma

## Abstract

Urinary bladder insult can be caused by environmental, genetic, and developmental factors. Depending upon insult severity, the bladder may lose its ability to maintain capacity and intravesical pressures resulting in renal deterioration. Bladder augmentation enterocystoplasty (BAE) is employed to increase bladder capacity to preserve renal function using autologous bowel tissue as a “patch.” To avoid the clinical complications associated with this procedure, we have engineered composite grafts comprised of autologous bone marrow mesenchymal stem cells (MSCs) with CD34+ hematopoietic stem/progenitor cells (HSPCs) co-seeded onto a pliable synthetic scaffold [POCO; poly(1,8-octamethylene-citrate-co-octanol)] or a biological scaffold (SIS; small intestinal submucosa) to regenerate bladder tissue in a baboon bladder augmentation model. We set out to determine the protein expression profile of bladder tissue that has undergone regeneration with the aforementioned stem cell seeded scaffolds along with baboons that underwent BAE. Data demonstrate that POCO and SIS grafted animals share high protein homogeneity between native and regenerated tissues while BAE animals displayed heterogenous protein expression between the tissues following long-term engraftment. We posit that stem cell seeded scaffolds can recapitulate tissue that is almost indistinguishable from native tissue at the protein level and may be used in lieu of procedures such as BAE.

**Significance of Study:** Bladder augmentation enterocystoplasty has been used for decades as the gold-standard surgical procedure to treat severely dysfunctional bladders. Unfortunately, bowel (typically ileum) used for augmentation is an anatomical and physiological mismatch to bladder tissue and results in numerous complications including bladder perforation, secondary and tertiary redo surgeries, metabolic imbalances, excess mucus production, and increased risk of cancer. This is in part due to the intestinal protein expression that serves as a starting point and subsequent foundation in the optimization of pseudo bladder tissue. Within the context of this study, we demonstrate that autologous, bone marrow derived mesenchymal stem cells along with primitive hematopoietic stem/progenitor cells can used to regenerate bladder tissue in a large deficit, non-human primate bladder augmentation model. Data demonstrate that this synergistic cellular combination facilitates the promotion of a protein tissue landscape that is nearly identical to native bladder tissue.

## 1 Introduction

Urinary bladder dysfunction can arise from neurologic conditions such as myelodysplasia, spina bifida (SB) exhibited in pediatric patients, spinal cord injury (SCI), traumatic brain injury (TBI), cerebral palsy, and military-based trauma. [1,2]. For combat military personnel, bladder injury is often a result of penetrating trauma [3]. Between October 2001 through January 2008, US military personnel who served overseas were found to exhibit bladder injury at rate of 21.3% (or 189) of the recorded 887 who had unique genitourinary (GU) injuries [4]. In a study by Kronstedt et al who analyzed the data set from the Department of Defense Trauma Registry (DoDTR) from January 1, 2007 to March 17, 2020, of the 2,584 GU combat injured soldiers who required surgery, 1,090 (42%) required bladder repairs. Additionally, in a study of 530 veterans enrolled in Trauma Infectious Disease Outcomes Study (TIDOS), 89 patients acquired genitourinary injuries from deployment within the period of 2009 – 2014. Of the 89 patients, 19 (21%) of the patients had bladder injury and a majority (52.6%) of these patients had UTIs [5]. The goal in managing a severely dysfunctional bladder includes preserving physiological function of the bladder so that urine can be stored under low pressure while maintaining its ability to void efficiently under volitional control. When conservative management fails, including the use of medicines or catheterization, surgical intervention in the form of urinary bladder diversion procedures or bladder augmentation enterocystoplasty (BAE) is often employed for the treatment of severe bladder conditions [6,7]. The ideal clinical outcome of these types of procedures is to increase bladder capacity and compliance by reducing intravesical pressure in order to protect renal function and improve on quality-of-life metrics [8].

BAE employs the use of autologous intestinal tissue including the ileum, gastric segments, or colonic segments. Unfortunately, this highly invasive surgical procedure poses unwanted long-term issues [1,6,8,9,10]. The ileum and colon promote the formation of bladder calculi at a rate of 3-52.5% and occurring as early as 5 months post-BAE [1,11,12]. The gastric segment is the last option and has a reduced risk of calculi but poses its unique complications such as hematuria-dysuria syndrome and increased risk of malignancy [9]. Additionally, other clinical issues arise over time including, excessive mucus production, electrolyte imbalances, and perforation can occur in all cases [13]. These issues are treatable in most cases but forces patients to perform intermittent self-catheterization to prevent bladder infection, calculi, and urinary tract infections. These issues can still arise even with ideal management, yet efficacy varies patient to patient. Although BAE is somewhat effective, the 10-year risk rate of re-do surgery is as high as 43.9% for spina bifida patients, for example [14].

Within the context of this study, we describe the proteomic profiling of three unique grafts used for bladder augmentation. These include the current gold standard for BAE (ileum), small intestininal submucosa (SIS) (a widely utilized biological scaffold) [15,16], and the highly reproducible synthetic scaffold, poly(1,8-octamethylene-citrate-co-octanol) (POCO) within the context of a non-primate (baboon) bladder augmentation model. Both the SIS and POCO scaffolds were co-seeded with autologous bone marrow derived mesenchymal stem cells (MSCs) with CD34+ hematopoietic stem/progenitor cells (HSPCs) prior to graft implantation [13,17,18]. Baboons underwent partial cystectomy and were then independently grafted with either autologous ileum in Enterocystoplasty (**E**), cell-seeded SIS (**CS-SIS**), or cell-seeded POCO (**CS-POCO**). At the conclusion of the study, native and regenerated (or ileum-augmented) bladder tissues were collected. The proteomic profiles of regenerated or ileum-augmented versus native bladder were analyzed and compared. This study is the first of its kind to demonstrate proteomic profiling in a large bladder tissue deficit, baboon bladder augmentation model that bears significant phylogenetic resemblance to human counterparts.

## 2 MATERIALS AND METHODS

### 2.1 Baboon bladder augmentation procedure

The study design for the bladder augmentation in the baboon (Papio anubis) model was described previously [12]. Briefly, a 50-65% bladder cystectomy was performed in the animal cohorts and the bladder deficit was augmented with either ileum (enterocystoplasty; **E**), cell-seeded (bone marrow derived, autologous MSCs and CD34+ HSPCs) biological scaffold (CS-SIS), and cell-seeded biodegradable and elastomeric scaffold (CS-POCO); n=3 baboons/group. Tissue-centric analyses utilized samples at 24M (**CS-POCO**); 24M (one animal at 27M, **E**); 26-29M (**CS-SIS**).

### 2.2 Protein and peptide purification

50 mg of tissue from each sample was homogenized in 1 ml lysis buffer containing 8 M urea, 1% SDS, in 50 mM HEPES pH 8.5, and HALT protease inhibitor cocktail (Thermo Fisher Scientific). The tissue extract was centrifuged at 3000g for 15 min to eliminate tissue debris and the supernatant was transferred to a new tube. 200 ug of protein from each sample was purified from impurities and lipids by methanol-chloroform precipitation and resuspended in 6M guanidine in 100 mM triethylammonium bicarbonate (TEAB). Proteins were reduced with 1 mM DTT and alkylated with 5 mM iodoacetamide, and were further diluted with 100 mM TEAB to minimize the guanidine hydrochloride concentration to less than 1.5 M before digestion with trypsin/lys-C protease mix, MS Grade, 1:50 ratio, (Thermo Fisher Scientific) overnight at 37°C. The digest was then acidified with formic acid to a pH of ∼2–4 and desalted by using C18 HyperSep cartridges. The purified peptide solution was dried and quantified using the Micro BCA Protein Assay Kit (Thermo Fisher Scientific). An equal amount of peptide (∼50 μg) from each sample was then used for isobaric tandem mass tag (TMT) labeling as per the manufacturer’s instructions (Thermo Fisher Scientific). After two hours of incubation at room temperature, the reaction was quenched with hydroxylamine at a final concentration of 0.3% (v/v). Isobaric-labeled samples were then combined and lyophilized. The combined isobaric labeled peptide samples were then fractionated by Pierce High pH Reversed-Phase Peptide Fractionation Kit to eight fractions per the manufacturer’s protocol. Fractions were then dried using a speed vac and reconstituted in LC-MS sample buffer (5% acetonitrile, 0.125% formic acid) until LC-MS/MS analysis and concentration were assessed using Micro BCA.

### 2.3 MS/MS tandem mass spectrometry

Purified peptides, 1.0 ug each, were loaded onto a Vanquish Neo UHPLC system (Thermo Fisher Scientific) with a heated trap and elute workflow with a c18 PrepMap, 5mm, 5μM trap column (P/N 160454) in a forward-flush configuration connected to a 25cm Easyspray analytical column (P/N ES802A rev2) 2μM, 100A, 75um x 25 with 100% Buffer A (0.1% formic acid in water) and the column oven operating at 35°C. Peptides were eluted over a 240 min gradient, using 80% acetonitrile, 0.1% formic acid (buffer B), starting from 2.5% to 10% over 10 min, to 30% over 140 min, to 40% over 60 min, to 65% over 18 min, then to 99% in 5 min and kept at 99% for 7 min, after which all peptides were eluted. Spectra were acquired with an Orbitrap Eclipse Tribrid mass spectrometer with FAIMS Pro interface (Thermo Fisher Scientific) running Tune 3.5 and Xcalibur 4.5 and using TMTpro SPS-MS3 RTS FAIMS method implemented in the Xcaliber software with Real Time search filter (RTS) for MS3 triggering. For all acquisition methods, spray voltage set to 1900V, and ion transfer tube temperature set at 300°C, FAIMS switched between CVs of −40 V, – 55 V, and −70 V with cycle times of 1.0s. MS1 detector set to orbitrap with 120 K resolution, wide quad isolation, mass range = normal, scan range = 400–1600 m/z, max injection time = 50 ms, AGC target = 300% microscans = 1, RF lens = 30%, without source fragmentation, and datatype = positive and centroid. Monoisotopic precursor selection was set to included charge states 2–6 and reject unassigned. Dynamic exclusion was allowed n = 1 exclusion for 40 s with 10ppm tolerance for high and low. An intensity threshold was set to 5000. Precursor selection decision = most intense. MS2 settings include isolation window = 0.7, scan range = auto normal, collision energy = 30% CID, scan rate = turbo, max injection time = 35 ms, AGC target = 1 × 104, Q = 0.25. In MS3, an on-line real-time search algorithm (Orbiter) was used to trigger MS3 scans for quantification. MS3 scans were collected in the Orbitrap using a resolution of 50,000, scan range 100-500, notch number =10, activation type HCD=55%, maximum injection time of 200 ms, and AGC of 200%. Isobaric tag loss exclusion was set to TMT pro reagent.

### 2.4 MS/MS data analysis

Raw data were analyzed using Proteome Discoverer 2.5 (Thermo Fisher Scientific) using the “PWF Tribrid TMTpro Quan SPS MS3 SequestHT Percolator” and “CWF Comprehensive Enhanced Annotation Reporter Quan” methods implemented in the PD2.5 software. The data were searched against the Baboon UniProt Protein Sequence Database [Papio anubis (species) Proteome Taxon ID 9555]. The search parameters included precursor mass tolerance of 10 ppm and 0.6 Da for fragments, allowing two missed trypsin cleavages, acetylation(+42.011 Da), Met-loss / −131.040 Da (M), and Met-loss+Acetyl (−89.030 Da (M) as N-terminal dynamic modification and carbamidomethylation (Cys), TMTpro / +304.207 Da in any N-terminus, TMTpro / +304.207 Da (K) as a static modification. Percolator PSM validation was used with the following parameters: strict false discover rate (FDR) of 0.01, relaxed FDR of 0.05, maximum ΔCn of 0.05, and validation based on q-value. Reporter ion quantitation was using the method TMTpro 18plexlotsWC320807, 18-plex Tandem Mass Tag® of Proteome Sciences plc method implemented on the proteome discoverer software and general quantification settings used with following settings, Peptides to Use: Unique + Razor; Consider Protein Groups for Peptide Uniqueness set as True; Normalization based on Total Peptide Amount; Scaling Mode set as none, low abundance peptides were removed by filtering out proteins with less than 3 PSMs, Protein ratio was calculated based on protein abundance and t-test were used for class comparison. Differentially expressed proteins were selected based on p-value < 0.05 and log_2_ fold change > 1.0 or < −1.0, or a 2-fold change. A positive fold change indicated a higher expression in the grafted or regenerated tissue compared to native tissue, while a negative fold change a higher expression in the native tissue compared to grafted or regenerated tissue. For uncharacterized proteins or proteins with unknown function presented in the manuscript, the UniProt Accession number was search against UniProt database https://www.uniprot.org/ and the NCBI Gene database https://www.ncbi.nlm.nih.gov/gene/. If required, more information was obtained with regards to protein identity by matching the amino acid sequence of the protein on the NCBI BLAST alignment program https://blast.ncbi.nlm.nih.gov/Blast.cgi.

### 2.5 Data normalization

Data was normalized by calculating the total sum of the abundance values for each channel over all the peptides identified. The channel with the highest total abundance was considered as a reference and correction was made for the abundance values in all other channels by a constant factor so that total abundance in all channels were the same.

## 3 RESULTS

### 3.1 Box plot distribution

In our study, we had examined the tissues collected from the native bladder tissue and compared against its own grafted tissue. There were three baboons in each of the three study groups, and 2 tissue samples from each animal, either grafted (for **E** group) or regenerated (for **CS-POCO** and **CS-SIS**) and native tissue samples, resulting in a total of 18 tissue samples as shown in Figure 1. The box plot was created to visualize the variation in abundances of mass spectrometric signals across different samples and conditions. As shown in the figure, the boxed area in each sample contained data that fell within Interquartile Range (IR), or 25-75% of the data range which includes data from Q1 to Q3, with the indicated median being at Q2. Any data points outside of the bars of the upper or lower region were single data points that are Q3 + 1.5IR or Q3 - 1.5IR, respectively. As shown in the figure, the IR for all 18 samples was similar and consistent across all samples indicating a uniform distribution of the data and suggested a tightly grouped result set. Since the median of the data was similar across all samples, there were minimal number of outliers and minimal variations variables or shifts amongst the samples.

**Figure 1:**
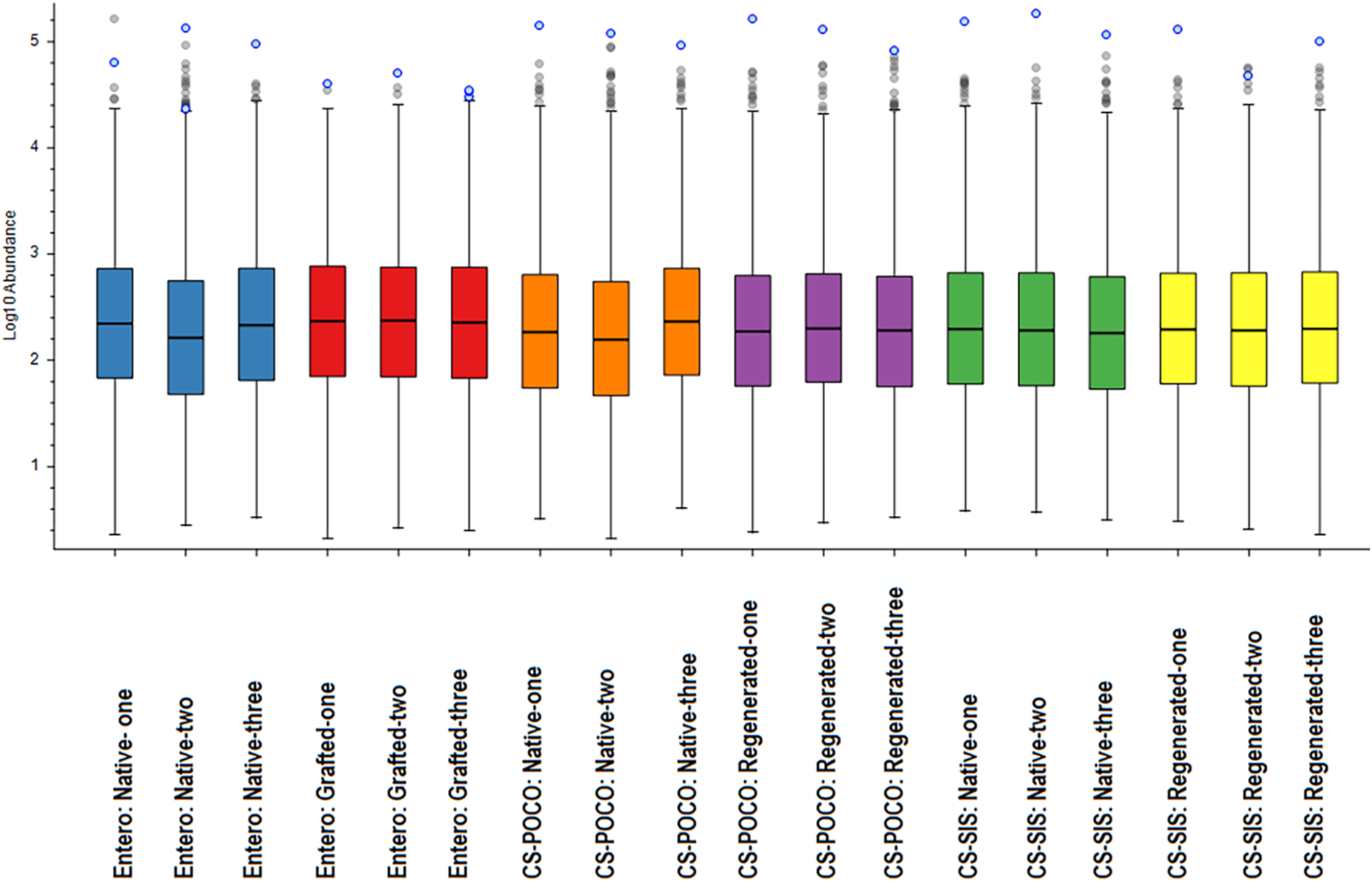
Box Plot Distribution of 18 tissue samples examined. For each study group-Entero, CS-POCO, and CS-SIS, there were 3 animals and for each animal the grafted or regenerated tissue was studied along with its matched native tissue sample. The colored box indicated the Interquartile Range of 25-75% of the data range with a line in the middle indicating the median. The outliers were single proteins shown for each of the sample.

### 3.2 Volcano plots

Proteomic profiles of the regenerated (or grafted) bladder tissue vs. the native bladder tissue for each animal were generated using our baboon bladder augmentation model. The data for the three animals were then averaged within each three groups (**E**, **CS-POCO,** and **CS-SIS**) of bladder augmentation. We analyzed a total of 5292 possible proteins and examined their expression patterns in the tissues within the aforementioned groups and determined the ratio of expression between the regenerated vs. native for each tissue graft type. The entire proteomic database for all 5292 proteins is included in the supplemental section, which contained expression levels for each tissue as well as differential expression levels for each regenerated/grafted bladder tissue type when compared to its own bladder native tissue.

For visualization of differential protein expression pattern, the proteomics data from the three groups was represented in our volcano plots, with the p-value graphed against the log_2_ fold ratio. Figures 2A, 2B, and 2C showed volcano plots of Entero: Grafted vs. Native, CS-POCO: Regenerated vs. Native, and CS-SIS: Regenerated vs. Native, respectively, comparing grafted tissue to its native tissue. For consideration for differential protein expression the p-value was less than 0.05 while the log_2_-fold change was either ≥ 1 or ≤ −1. As indicated on the volcano plots, the areas where the proteins that were indicated as differentially expressed had log_10_ p-value > 1.3 (or p-value < 0.05), and log_2_ fold ratio ≥ 1 for the proteins expression in the grafted or regenerated tissue to be higher than its native tissue (as marked by pink box) or log_2_ fold ratio ≤ −1 for protein expression in the grafted or regenerated tissue to be lower than its native tissue (marked in blue box). As shown in Figure 2A the **E** group with the autograft ileum, when comparing the grafted tissue to the native tissue, of the 5292 proteins that were surveyed, 160 proteins were expressed at higher level in the native tissue compared to the **E** grafted tissue at a log_2_ fold ratio ≤ −1, both at p-value < 0.05. The information of these 160 proteins were included in Table 1A. Conversely, 259 proteins had higher expression in the E grafted tissue than the native tissue at a log_2_ fold ratio ≥ 1 and are detailed in Table 1B. In Figure 2B, the **CS-POCO** study group, no protein in the generated tissue had significant overexpression over the native tissue, while 2 proteins had log_2_ fold ratio ≤ −1 or higher expression level in native tissue than **CS-SIS** regenerated tissue, at significant level of p-value < 0.05. In Figure 2C, the **CS-SIS** study group, there was 1 protein that had a log_2_ fold ratio ≥ 1 (expressed in higher levels in **CS-SIS** regenerated tissue than its native tissue) and 5 proteins that had log_2_ fold ratio ≤ −1 (expressed at higher level native protein than **CS-SIS** regenerated tissue), also all had a p-value < 0.05. The differently expressed proteins are shown in the tables in the following section, where Table 1A and Table 1B include differentially expressed proteins in the E group as represented in Figure 2A, while Table 1C and Table 1D correspond to Figure 2B and Figure 2C, respectively.

**Figure 2:**
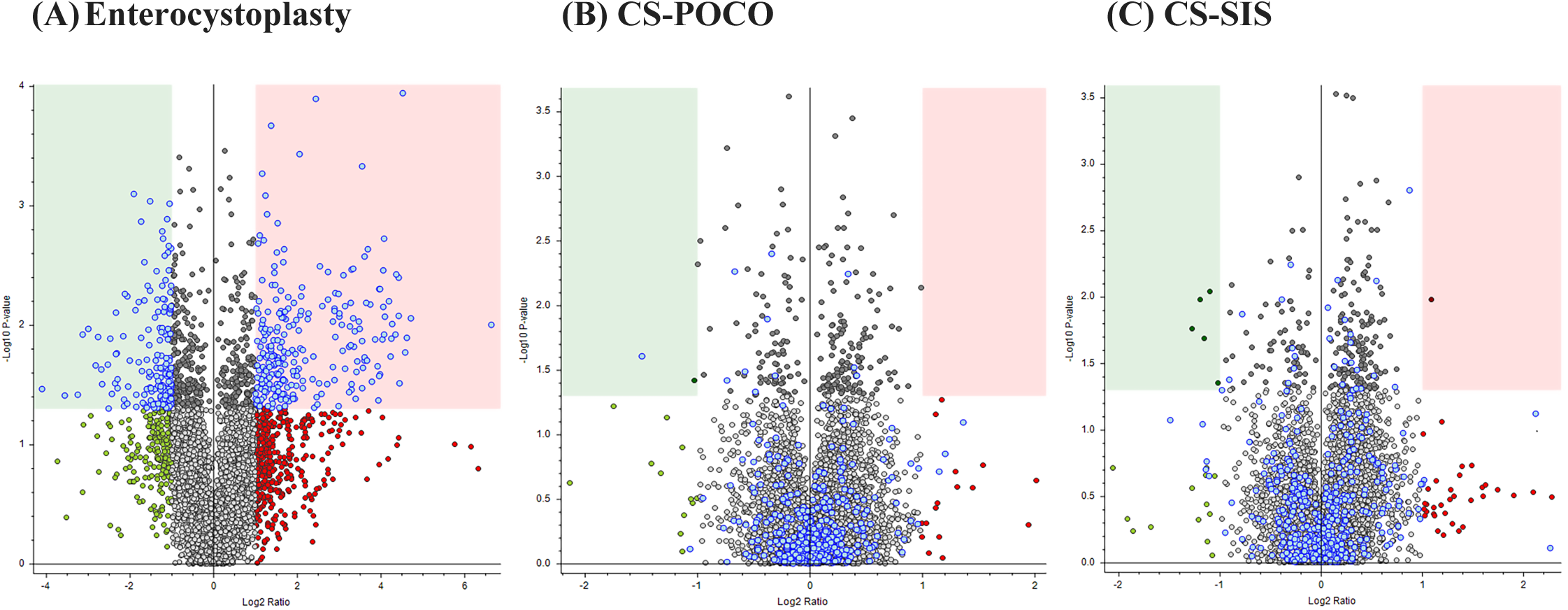
Volcano Plots showing the differentially expressed protein distribution of the grafted or regenerated tissue vs. the native tissue. Each protein is shown as a dot in the graph. Proteins in the green colored box indicated proteins that were highly expressed in the native tissue vx. the grafted or regenerated tissue with log2 fold ratio of < −1 and p-value < 0.05. Proteins in the pink colored box indicated proteins that were highly expressed in the grafted or regenerated tissue vs. the tissue with log2 fold ratio of > 1 and p-value < 0.05. (A) Entero: Grafted vs. Native tissue showed many differentially expressed proteins in both grafted and native tissue. (B) CS-POCO: Regenerated vs. Native tissue showed the least number of proteins differentially expressed in either tissue. (C) CS-SIS: Regenerated vs. Native showed 4 proteins highly expressed in the Native tissue and 1 protein in the Regenerated tissue.

**Table lA.**
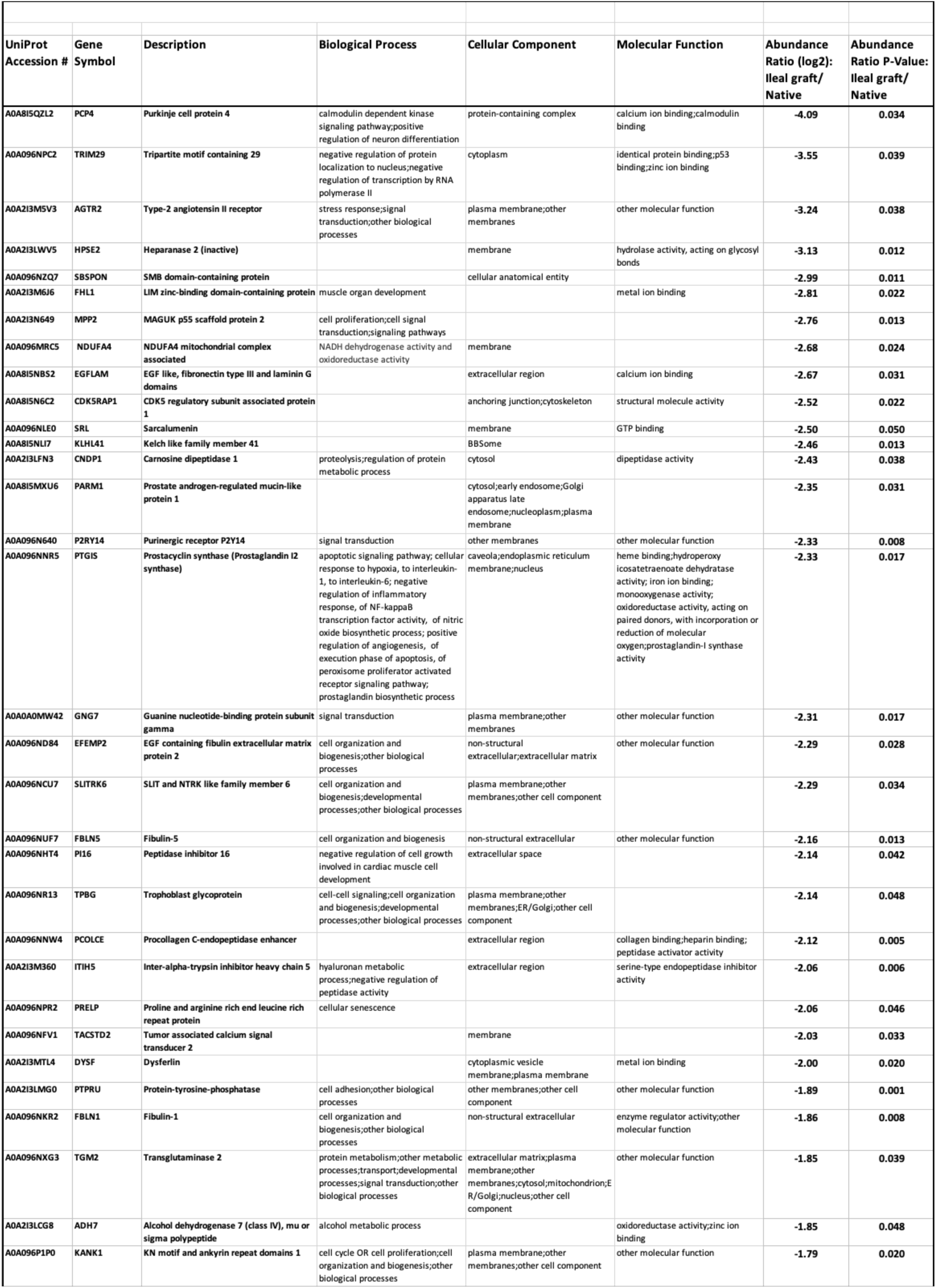
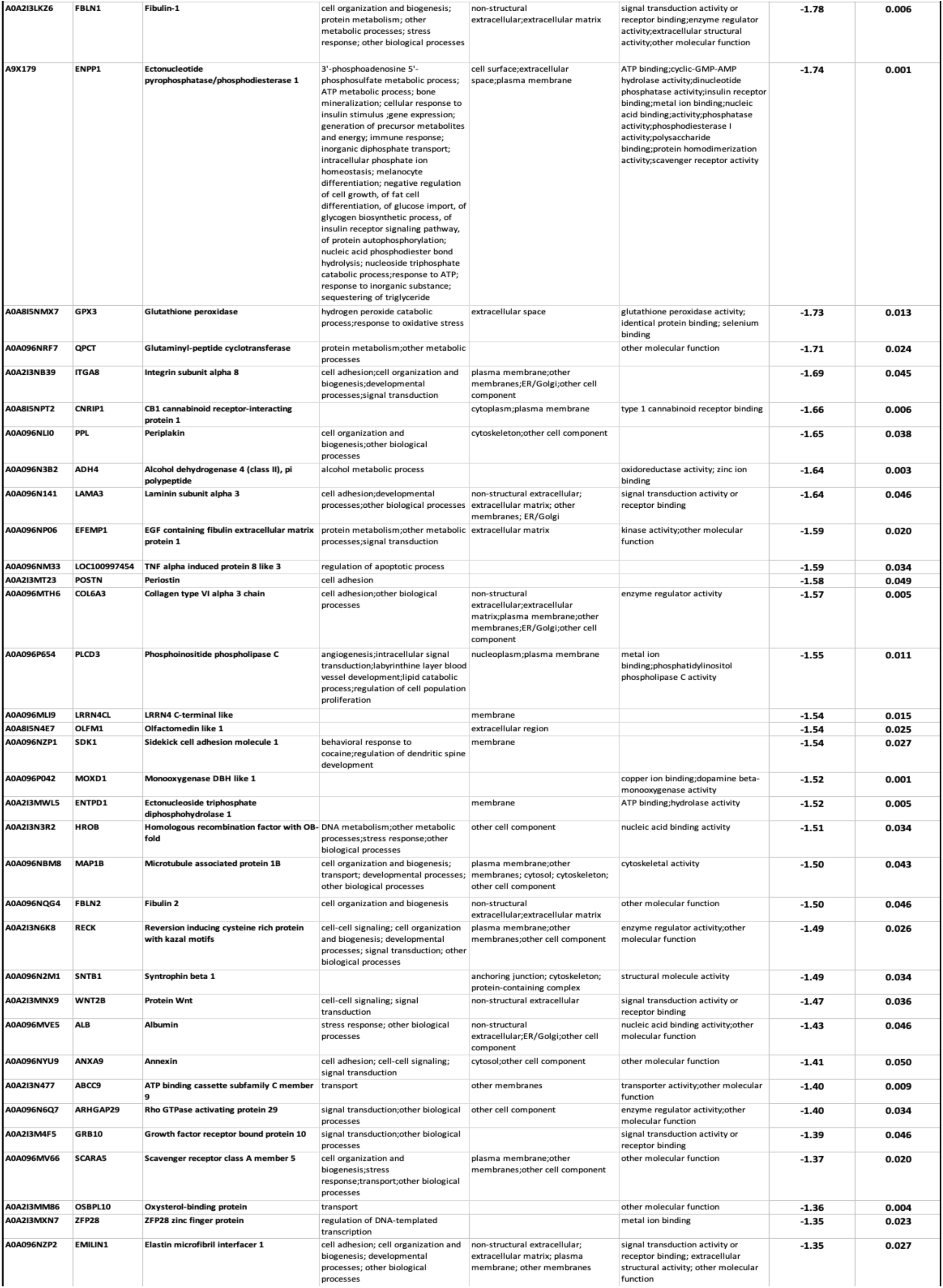
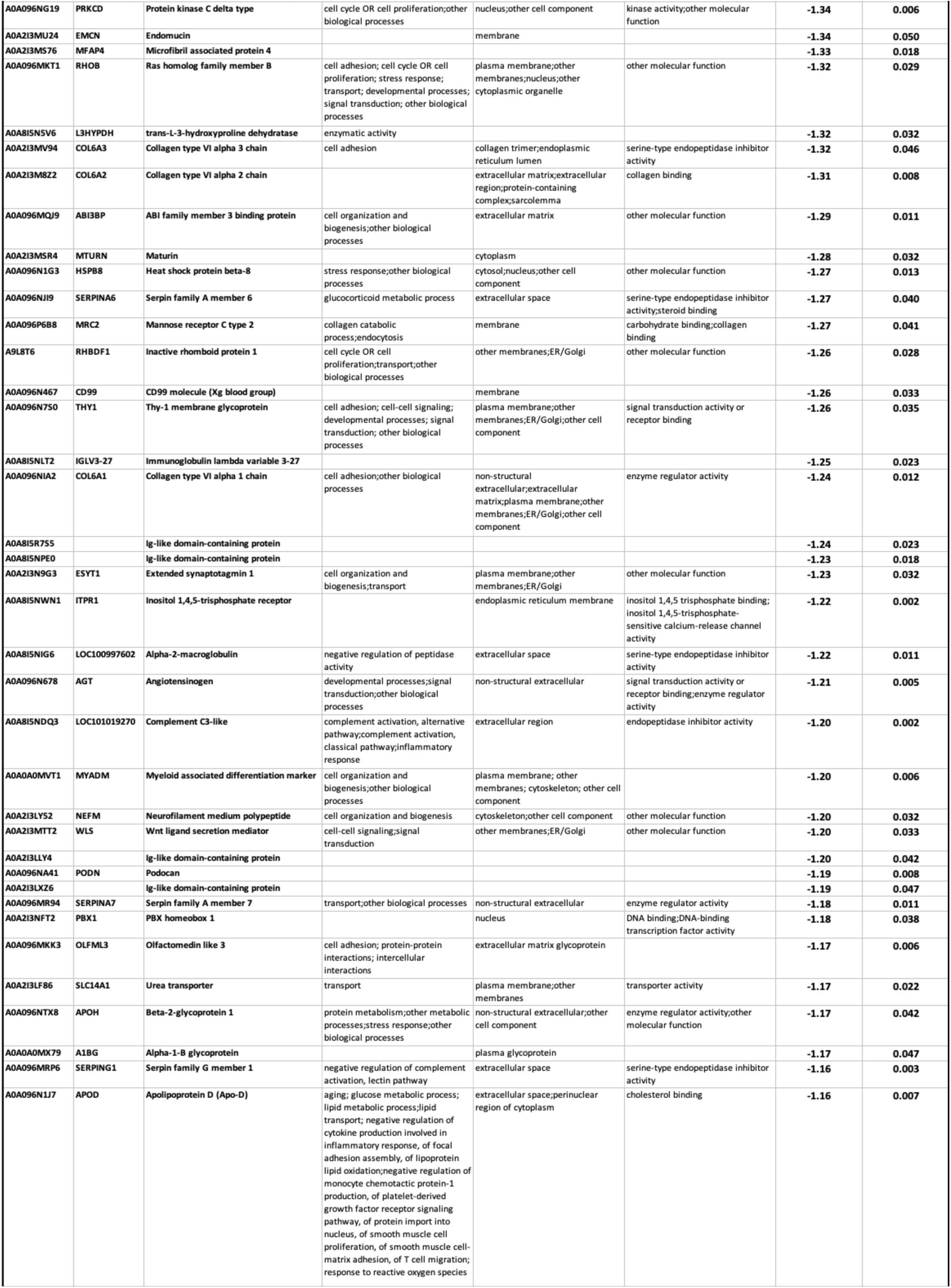
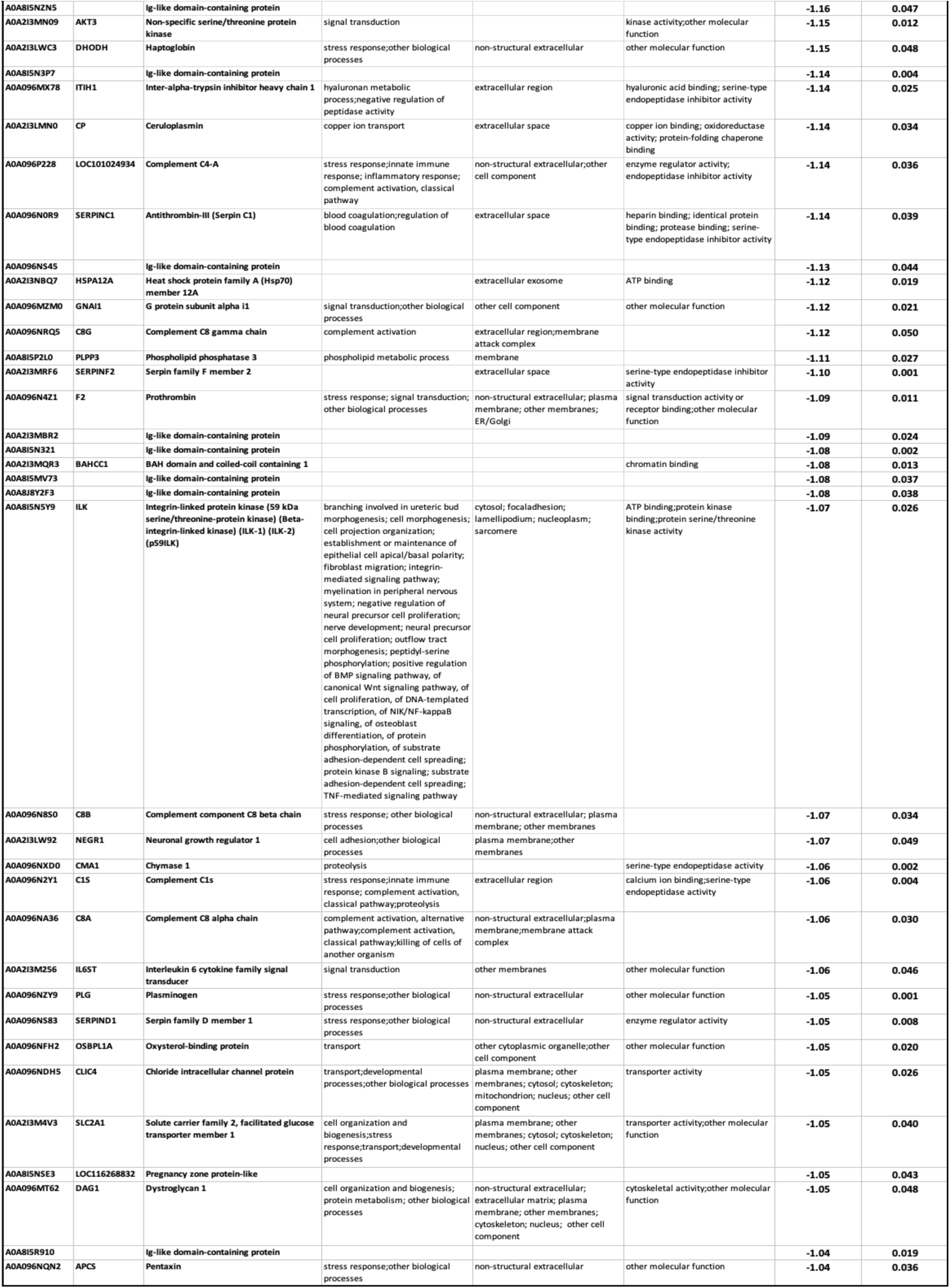
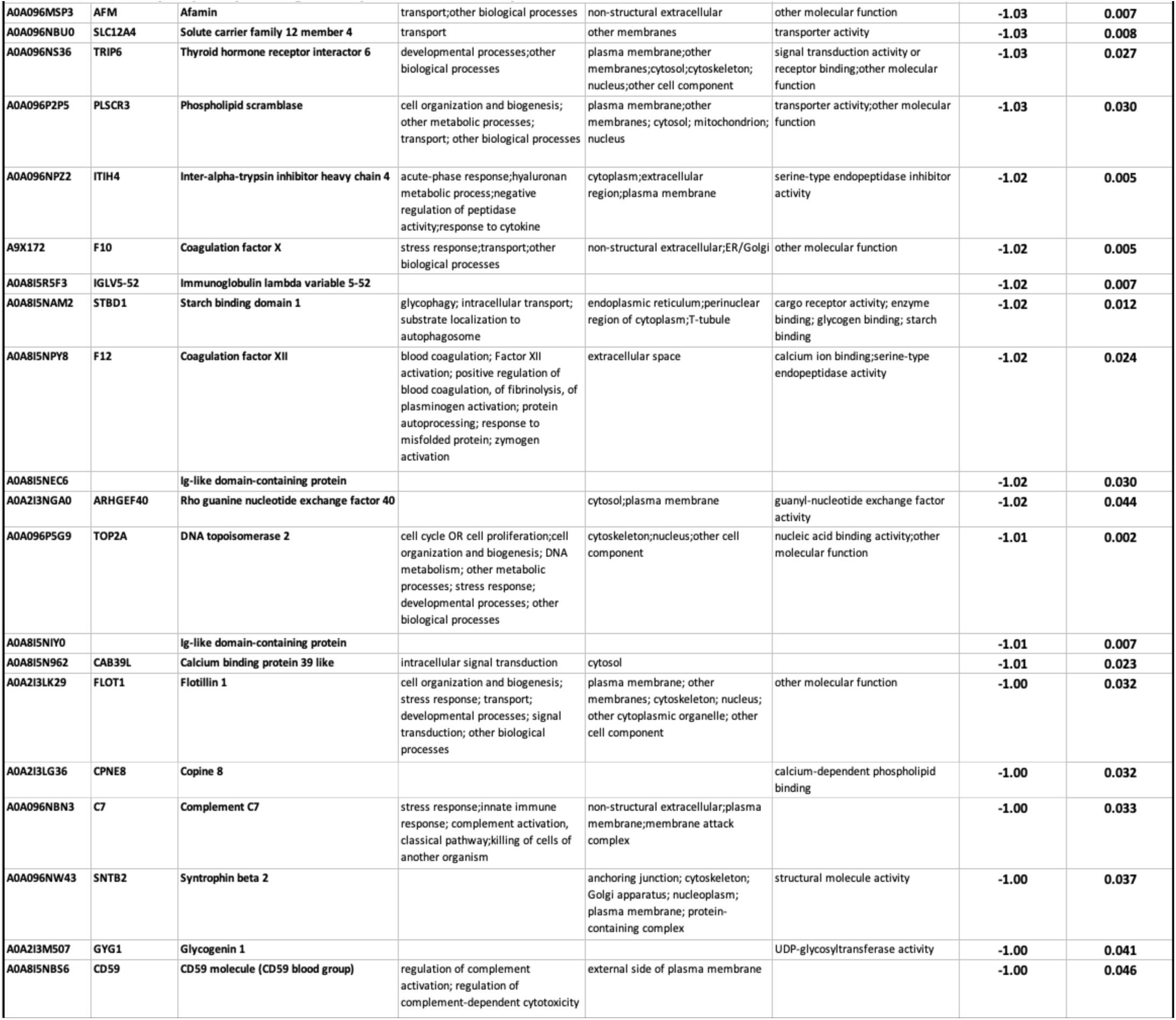
Enterocystoplasty-Ileal grafted proteins < Native proteins.

**Table 1B.**
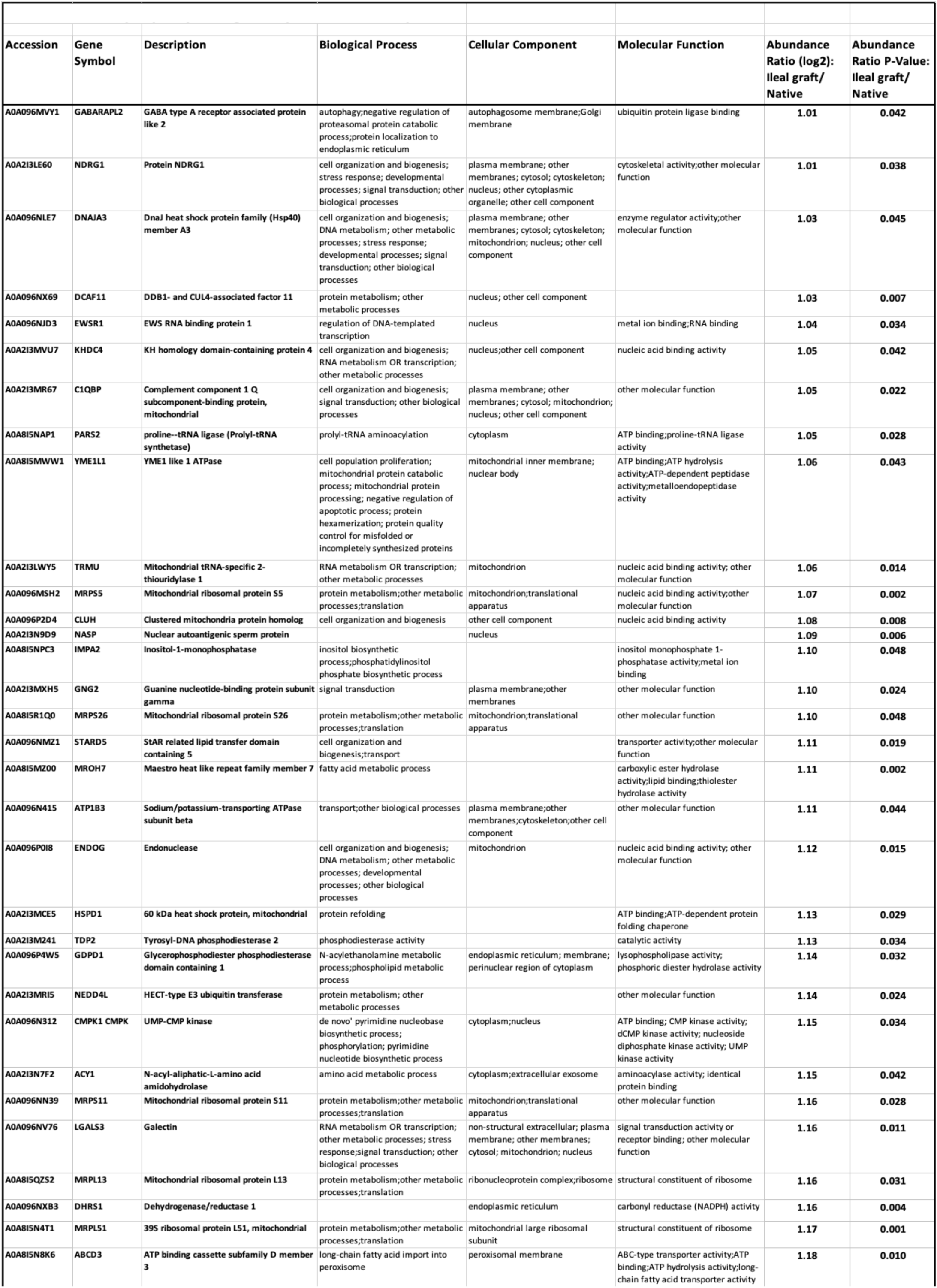
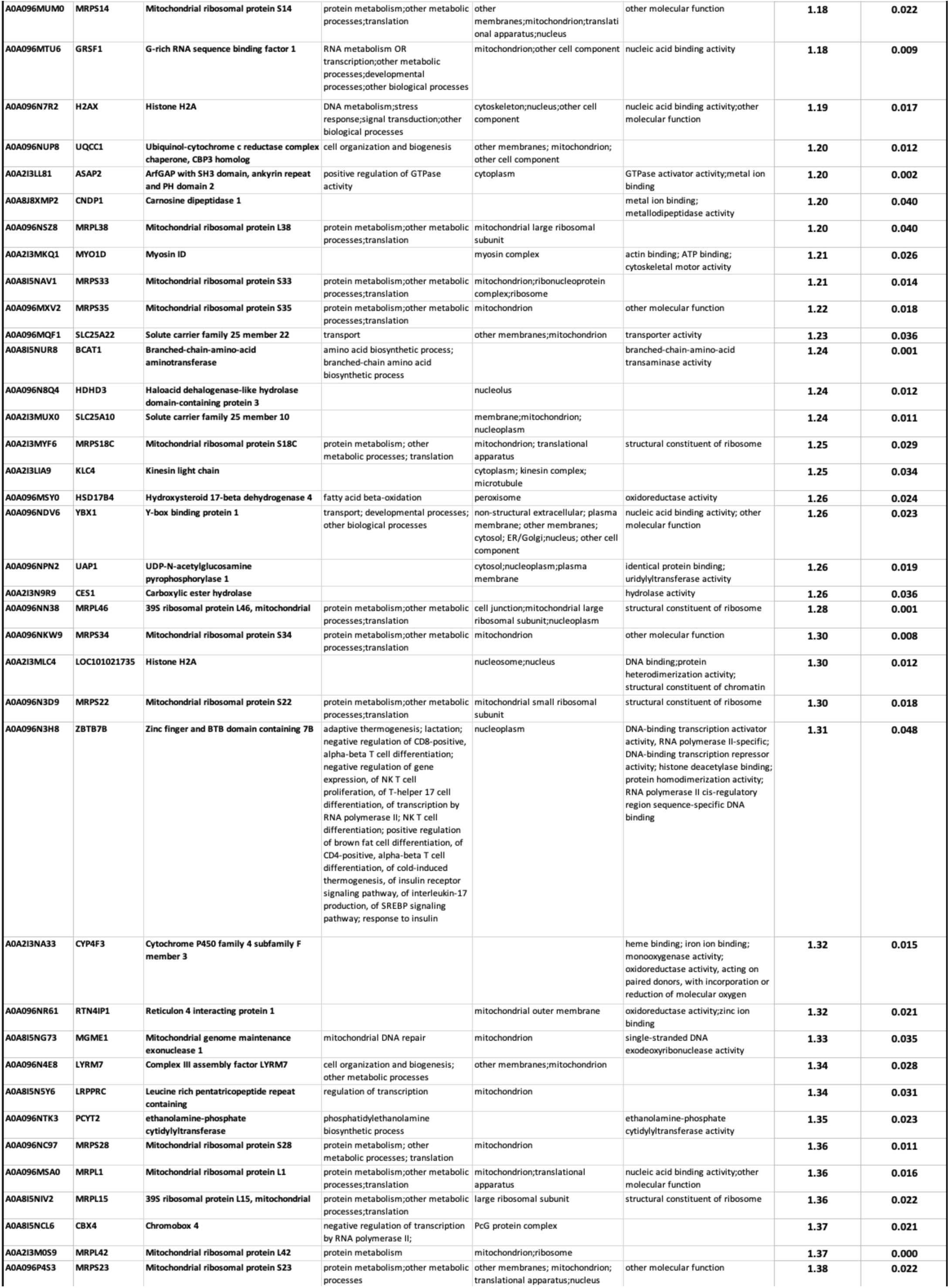
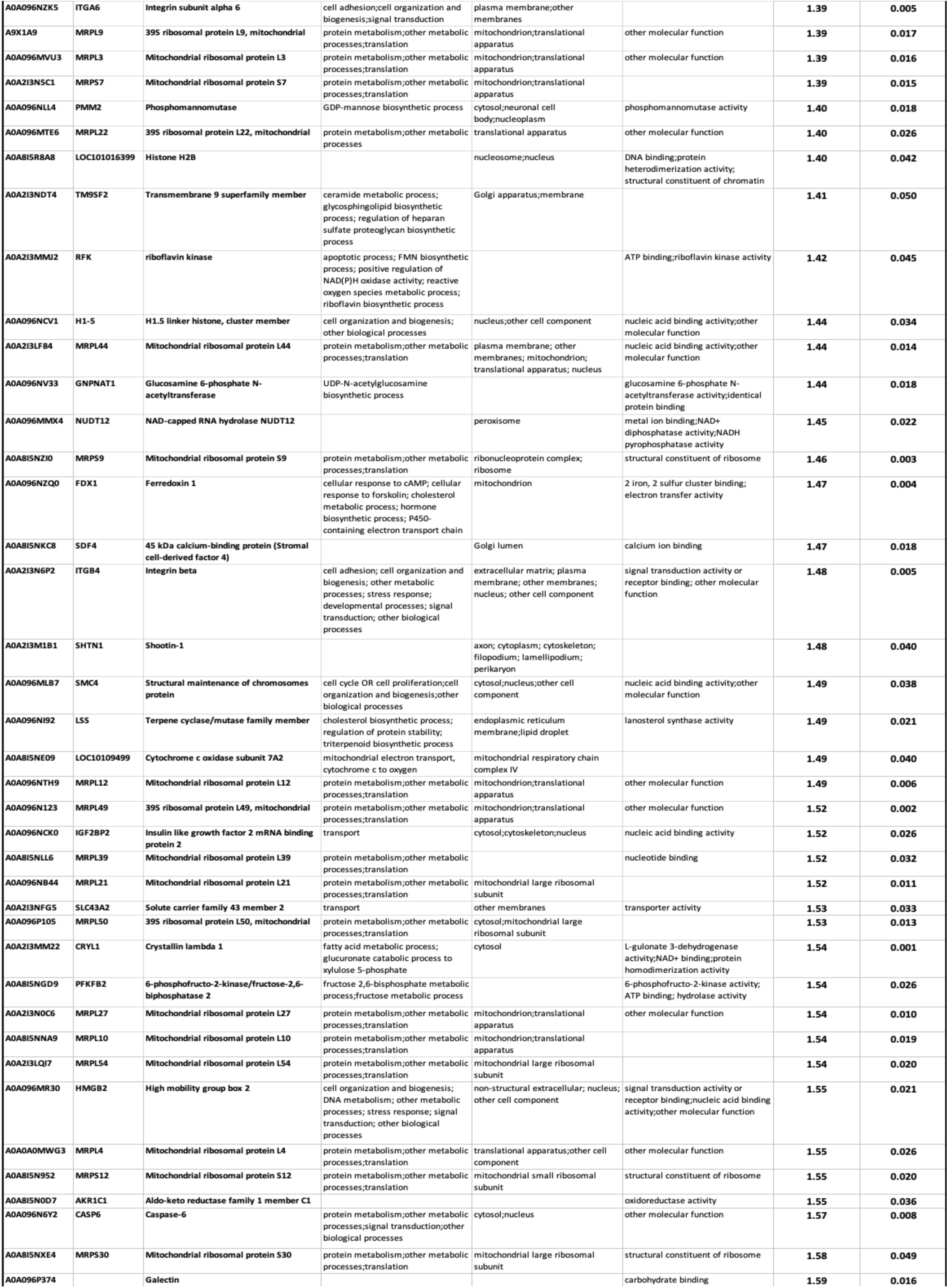
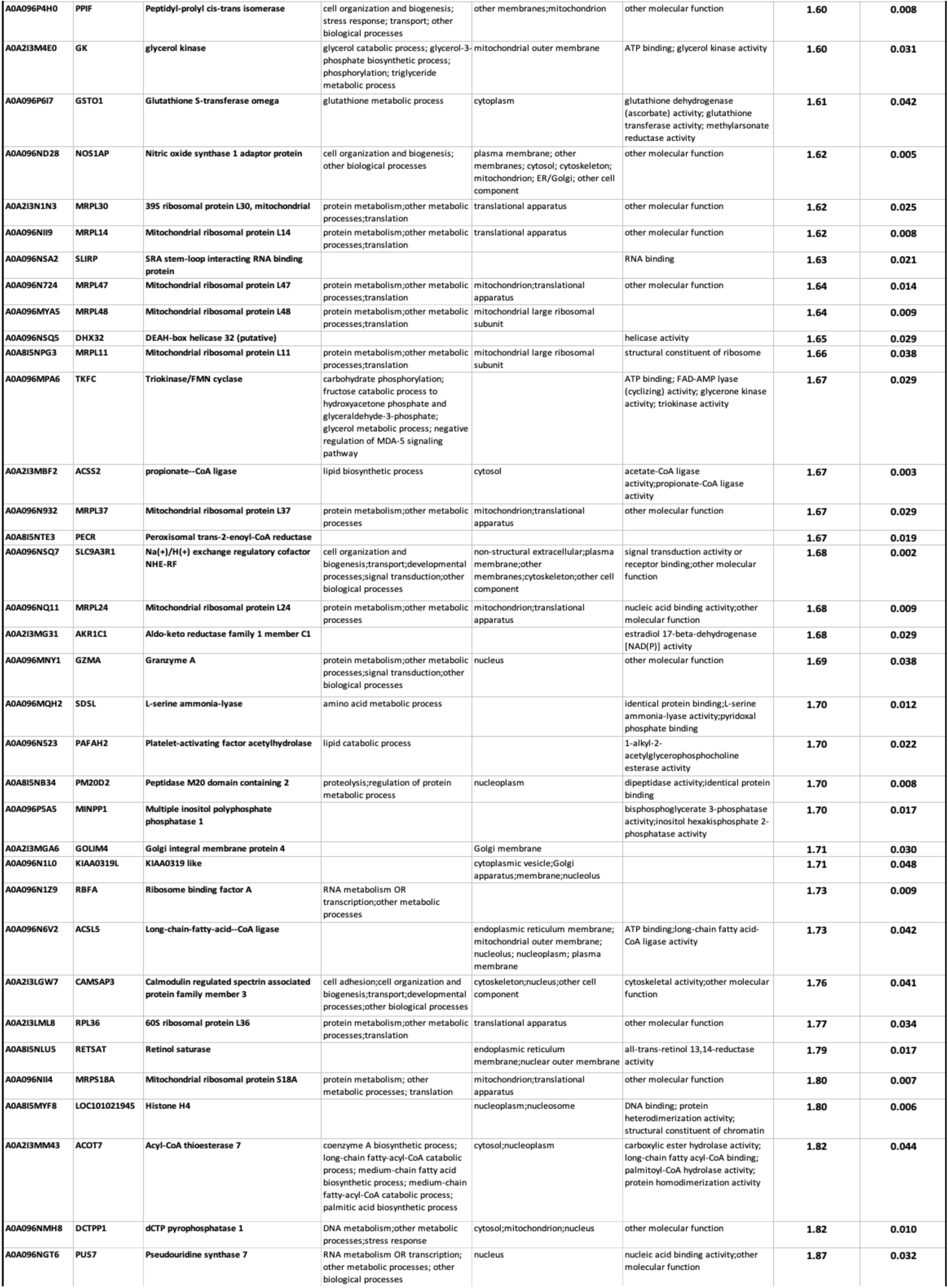
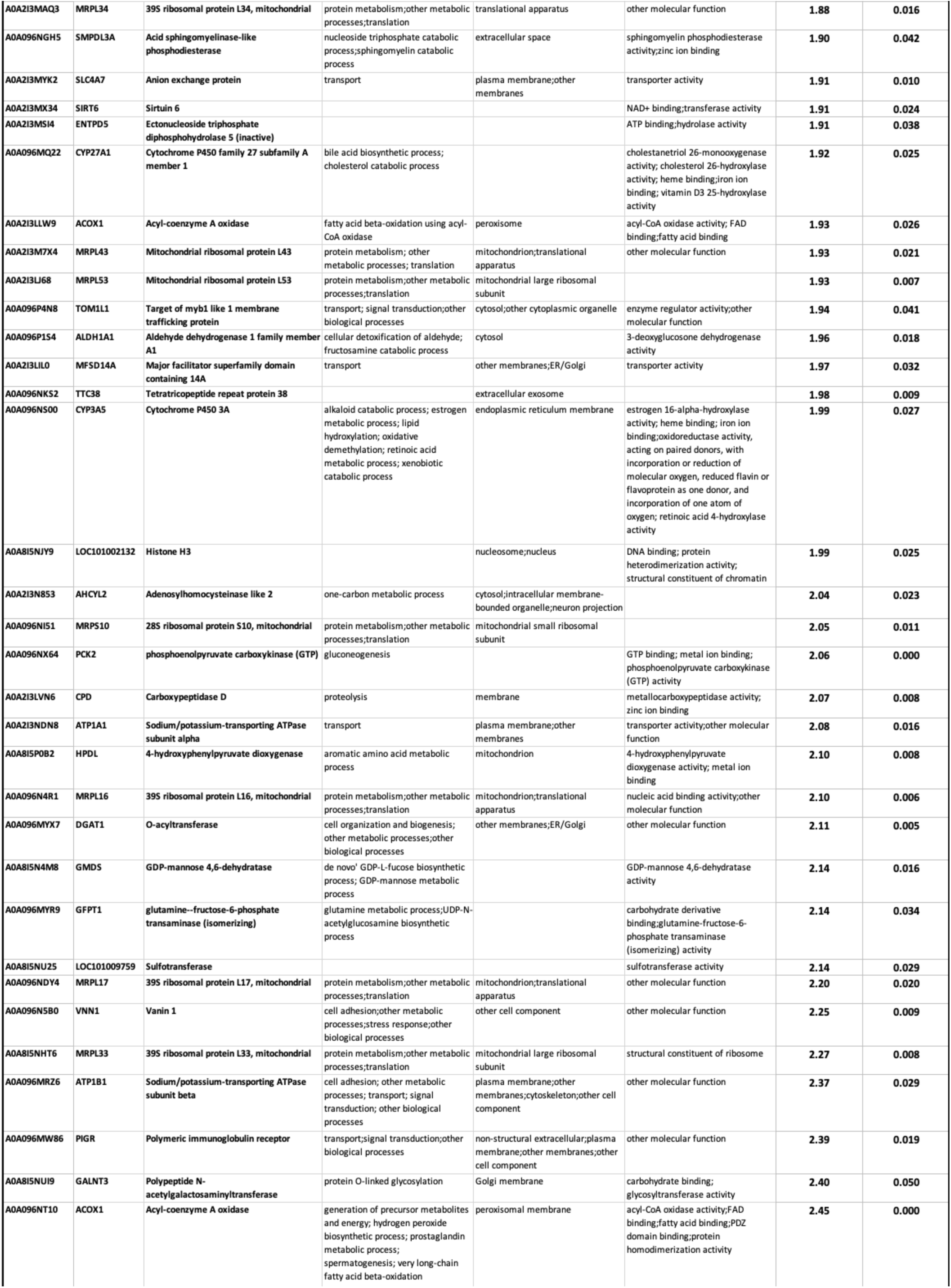
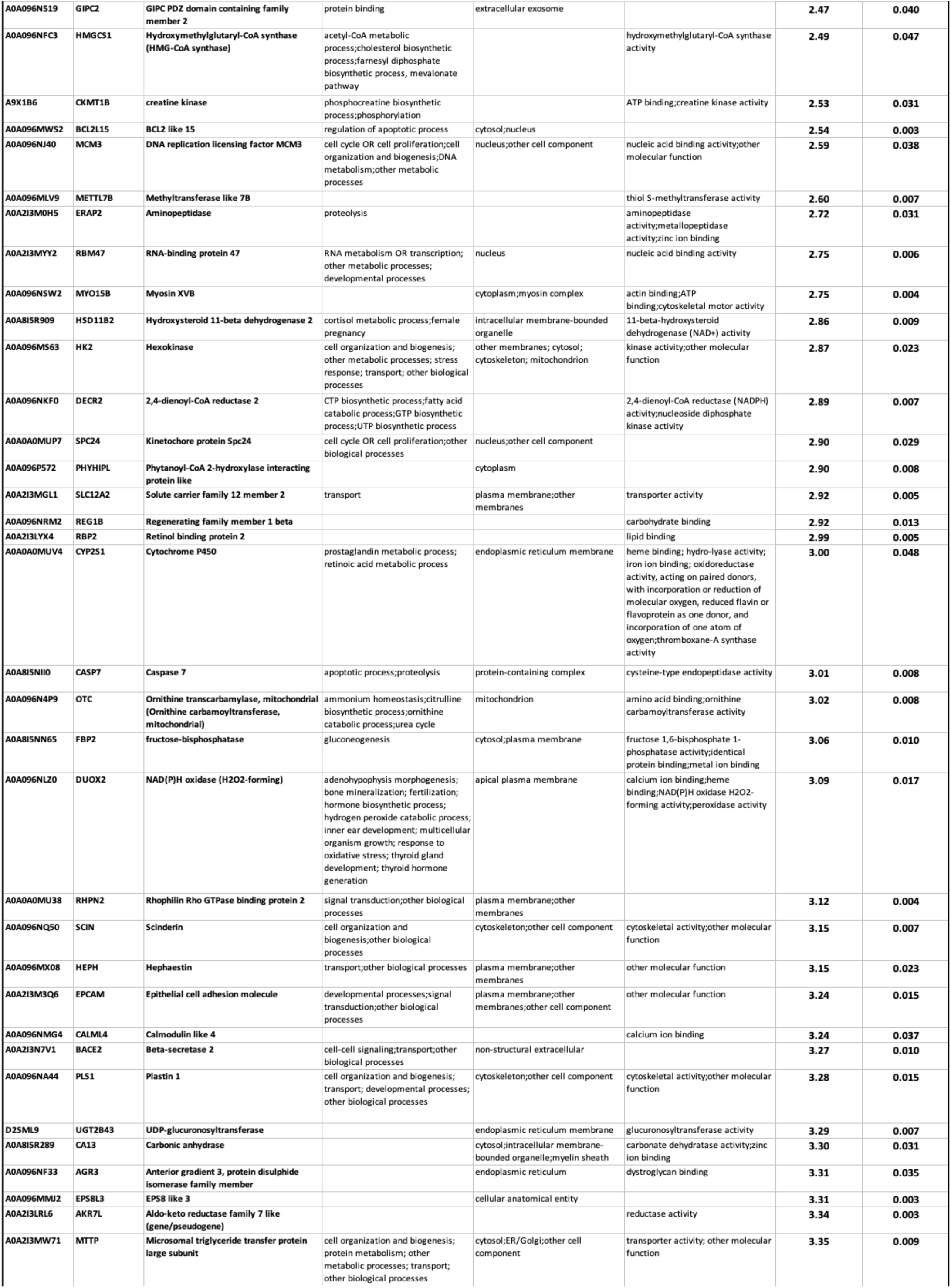
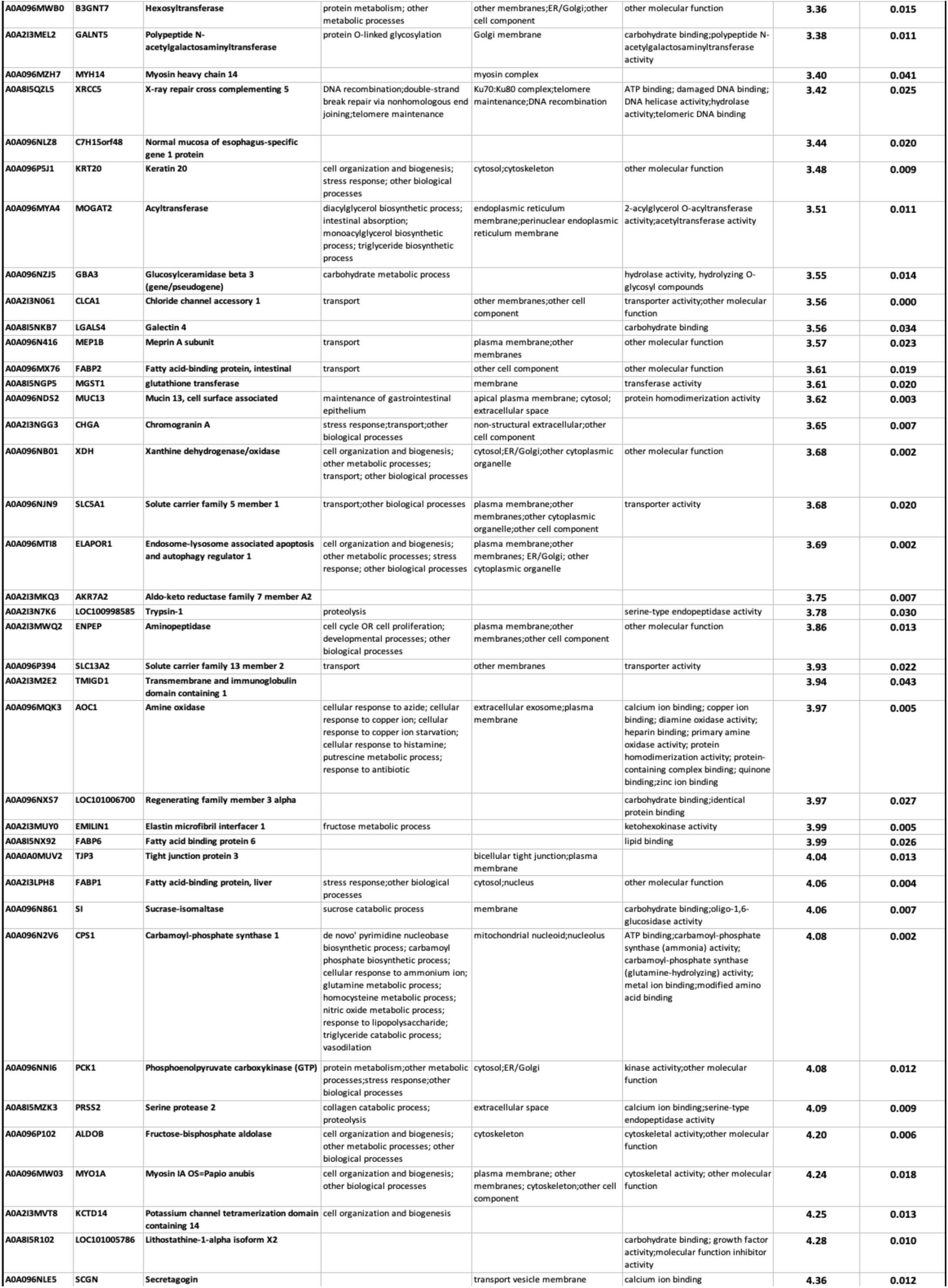
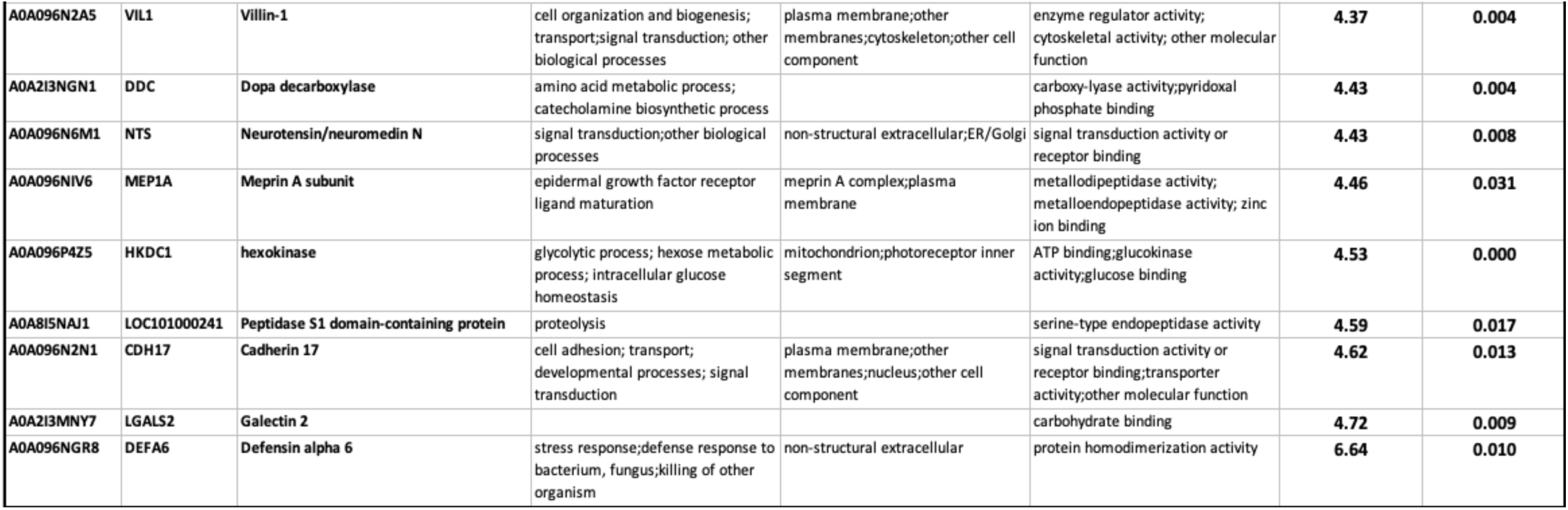
Enterocystoplasty-Ileal grafted proteins < Native proteins.

**Table 1C.** CS-POCO-Regenerated proteins < Native proteins.

| Table 1C. CS-POCO- Regenerated proteins < Native proteins |  |  |  |  |  |  |  |
| --- | --- | --- | --- | --- | --- | --- | --- |
| Accession | Gene Symbol | Description | Biological Process | Cellular Component | Molecular Function | Abundance Ratio (log2): Regenerated/ Native | Abundance Ratio P-Value: Regenerated/ Native |
| A0A096NFG4 | CHRD1 | Chordin like 1 |  |  | developmental protein | -1.030 | 0.038 |
| A0A2I3MNY7 | LGALS2 | Galectin-2 | carbohydrate binding |  | other molecular function | -1.490 | 0.025 |

**Table 1D.**
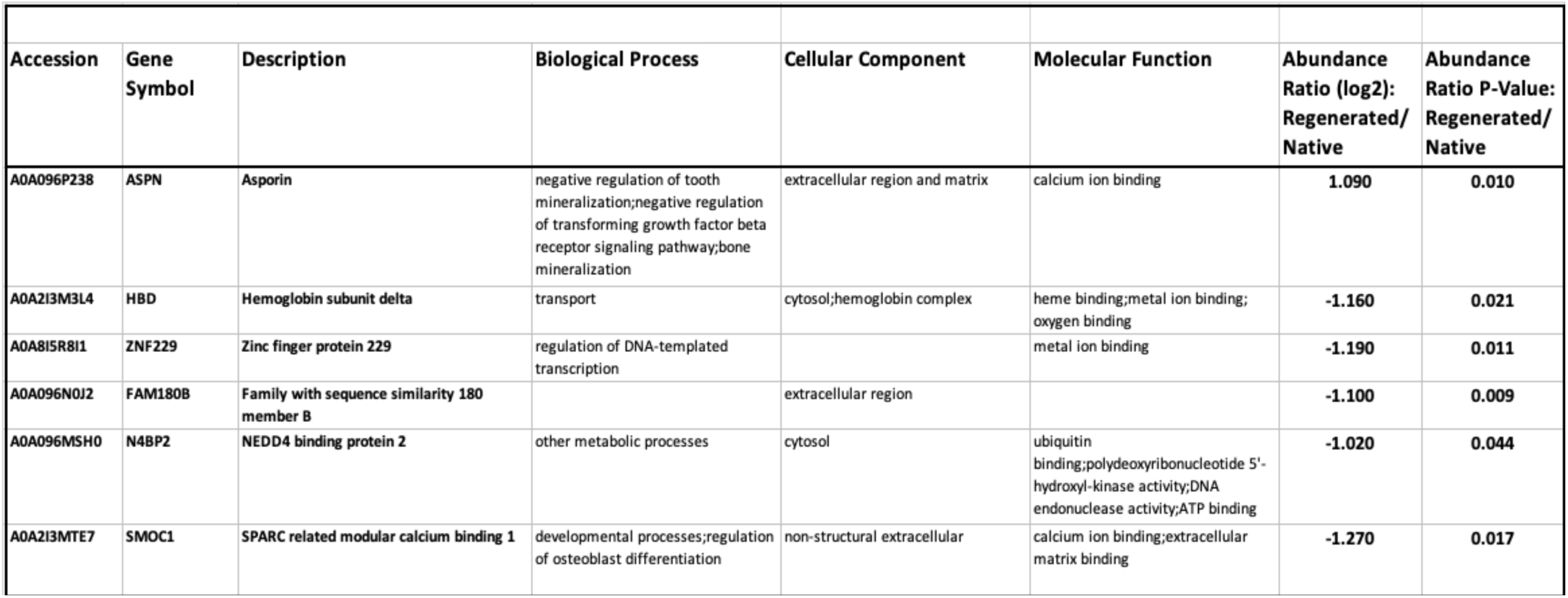
CS-SIS-Regenerated proteins vs. Native proteins.

| Table 1D. CS-SIS- Regenerated proteins vs. Native proteins |  |  |  |  |  |  |  |
| --- | --- | --- | --- | --- | --- | --- | --- |
| Accession | Gene Symbol | Description | Biological Process | Cellular Component | Molecular Function | Abundance Ratio (log2): Regenerated/ Native | Abundance Ratio P-Value: Regenerated/ Native |
| A0A096P238 | ASPN | Asporin | negative regulation of tooth mineralization; negative regulation of transforming growth factor beta receptor signaling pathway; bone mineralization | extracellular region and matrix | calcium ion binding | 1.090 | 0.010 |
| A0A2I3M3L4 | HBD | Hemoglobin subunit delta | transport | cytosol; hemoglobin complex | heme binding; metal ion binding; oxygen binding | -1.160 | 0.021 |
| A0A8ISR8I1 | ZNF229 | Zinc finger protein 229 | regulation of DNA-templated transcription |  | metal ion binding | -1.190 | 0.011 |
| A0A096N0J2 | FAM180B | Family with sequence similarity 180 member B |  | extracellular region |  | -1.100 | 0.009 |
| A0A096MSH0 | N4BP2 | NEDD4 binding protein 2 | other metabolic processes | cytosol | ubiquitin binding; polydeoxyribonucleotide 5'-hydroxyl-kinase activity; DNA endonuclease activity; ATP binding | -1.020 | 0.044 |
| A0A2I3MTE7 | SMOC1 | SPARC related modular calcium binding 1 | developmental processes; regulation of osteoblast differentiation | non-structural extracellular | calcium ion binding; extracellular matrix binding | -1.270 | 0.017 |

### 3.3 Differentially expressed proteins

For the consideration of differentially expressed protein, the p-value for the protein was lower than 0.05 (a 95% confidence interval). After setting this threshold, we examined all the proteins that were highly expressed in the grafted or regenerated tissue or its matched native tissue by comparing their abundance within those tissues by their log_2_ fold ratio. The negative log_2_ fold ratio designates a protein with higher expression level in the native tissue when compared to the regenerated grafted tissue. The differentially expressed proteins are shown in Table 1A for **E** grafted proteins < native proteins, Table 1B for **E** grafted proteins > native proteins, Table 1C for **CS-POCO**, and Table 1D for **CS-SIS**. For each table, the identifying feature of the protein is the UniProt Accession number. Any uncharacterized proteins or proteins with unknown function were updated using information listed on the UniProt and GenBank Gene websites.

As shown in Table 1C, for the **CS-POCO** graft, only two proteins were significantly differentially expressed. CHRDL1 (chordin like 1) had log2 fold ratio of −1.03 (p-value= 0.03836) while LGALS2 (Galectin-2) had a log_2_ fold ratio of −1.49 (p-value= 0.02488). Both proteins had higher expression in the native tissue when compared to the **CS-POCO** graft regenerated tissue. For Table 1D, protein ASPN (asporin), which had an expression value of log_2_ fold ratio of 1.09, while 5 proteins, Hemoglobin, ZNF229 (Zinc finger protein 229), FAM180B (Family with sequence similarity 180 member B), N4BP2 (NEDD4 binding protein 2), SMOC1 (SPARC related modular calcium binding 1), all were expressed at higher expression in the native tissue compared to the CS-SIS tissue at log_2_ fold ratio ranging from −1.27 to −1.02.

Tables 1A and 1B showed the differentially expressed proteins for the **E** group which had the greatest number of proteins that differed significantly between the native tissue and the regenerated tissue. As shown, within the 419 proteins that were differentially expressed, of which 160 proteins were expressed at higher levels in the native tissue compared to regenerated tissue in Table 1A, while 259 proteins with higher expression levels in the entero grafted tissue compared to the native tissue in Table 1B. In Table 1A, the top 10 proteins having the highest abundance ratio in the native tissue compared to entero grafted tissue included Purkinje cell protein 4 (PCP4) (abundance log_2_ fold ratio of −4.09, p-value of 0.034), Tripartite motif containing 29 (TRIM29) (abundance log2 fold ratio of −3.55, p-value of 0.039), Type-2 angiotensin II receptor (AGT2) (abundance log2 fold ratio of −3.24, p-value of 0.038), Heparanase 2 (inactive)-HPSE2 (abundance log_2_ fold ratio of −3.13, p-value of 0.012), SMB domain-containing protein (SBSPON) (abundance log2 fold ratio of −2.99, p-value of 0.011), LIM zinc-binding domain-containing protein (FHL1) (abundance log2 fold ratio of −2.81, p-value of 0.022), MAGUK p55 scaffold protein 2 (MPP2) (abundance log2 fold ratio of −2.76, p-value of 0.013), uncharacterized protein (abundance log2 fold ratio of −2.68, p-value of. 0.024), EGF like, fibronectin type III and laminin G domains (EGFLAM) (abundance log2 fold ratio of −2.67, p-value of 0.031), and CDK5 regulatory subunit associated protein 1 (CDK5RAP1) (abundance log2 fold ratio of −2.52, p-value of 0.022).

In Table 1B, the top 10 proteins with the highest abundance ratio in the **E** grafted tissue compared to native tissue included Defensin alpha 6 (DEFA6) (abundance log_2_ fold ratio of 6.64, p-value of 0.010), Galectin 2 (LGALS2), (abundance log_2_ fold ratio of 4.72, p-value of 0.009), Cadherin 17 (CDH17) (abundance log_2_ fold ratio of 4.62, p-value of 0.013), Peptidase S1 domain-containing protein (LOC101000241) (abundance log_2_ fold ratio of 4.59, p-value of 0.017), Hexokinase-HKDC1 (abundance log_2_ fold ratio of 4.53, p-value of 0.000), Meprin A subunit (MEP1A) (abundance log_2_ fold ratio of 4.46, p-value of 0.031), Neurotensin/neuromedin N (NTS) (abundance log_2_ fold ratio of 4.43, p-value of 0.008), Dopa decarboxylase (DDC) (abundance log_2_ fold ratio of 4.43, p-value of 0.004), Villin-1 (VIL1) (abundance log_2_ fold ratio of 4.37, p-value of 0.004), and Secretagogin (SCGN) (abundance log_2_ fold ratio of 4.37, p-value of 0.004).

The biological function of the proteins was categorized as illustrated in Figure 3 with regards to those proteins expressed in **E** and summarized in Tables 1A and 1B. The gene symbol, protein description, biological function, cellular component, and molecular function were updated when needed to the most up to date UniProt and UniGene databases. As shown in the figure, the distribution of proteins that were highly expressed in the grafted tissue was indicated in the blue color while the native tissue in orange color. The protein categories based on biological process included amino acid metabolism, apoptotic process, ATP binding, calcium binding, carbohydrate binding/process, cell adhesion, cell cycle or cell proliferation, cell-cell signaling, cell organization and biogenesis, cholesterol process, complement activity, DNA binding/chromatin binding/or RNA binding, DNA metabolism, fatty acid metabolism/processing/lipid binding, fructose/mannose/or glycerol process/glucogenesis/glycolytic process, immunoglobulin/Ig-like proteins, metal ion/copper/zinc/or iron ion binding, protein metabolism/proteolysis, serine-type endopeptidase inhibitor activity, signal transduction, stress response, structural activity, transport, and other biological process/cellular component/or nuclear function. Some of the proteins that had overlapping functions were categorized according to the first listed function. In the figure for the 259 proteins that were highly expressed at levels greater than log2-fold ratio in the **E**-grafted tissue when compared to the native tissue are shown in blue, while the 160 proteins at highly in the native tissue vs. **E**-grafted tissue are shown in orange. As shown in the distribution graph, in several categories, more proteins were highly expressed in one tissue vs the other tissue, or none in the other tissue. An example of this included the category of amino acid metabolism where the **E**-grafted tissue had 7 proteins that are highly expressed and none in the native tissue. For proteins that have complement activity, there were 6 proteins that were highly expressed in the native tissue vs. none for the **E**-grafted tissue. This scenario is similar for the serine-type endopeptidase inhibitor activity category with 9 proteins in the native vs. none in the **E**-grafted tissue. For the protein metabolism/ proteolysis category, a large number of proteins-63 were highly expresses in the **E**-grafted tissue compared to native, while 6 were highly expressed in the native tissue compared to **E**-grafted tissue.

**Figure 3:**
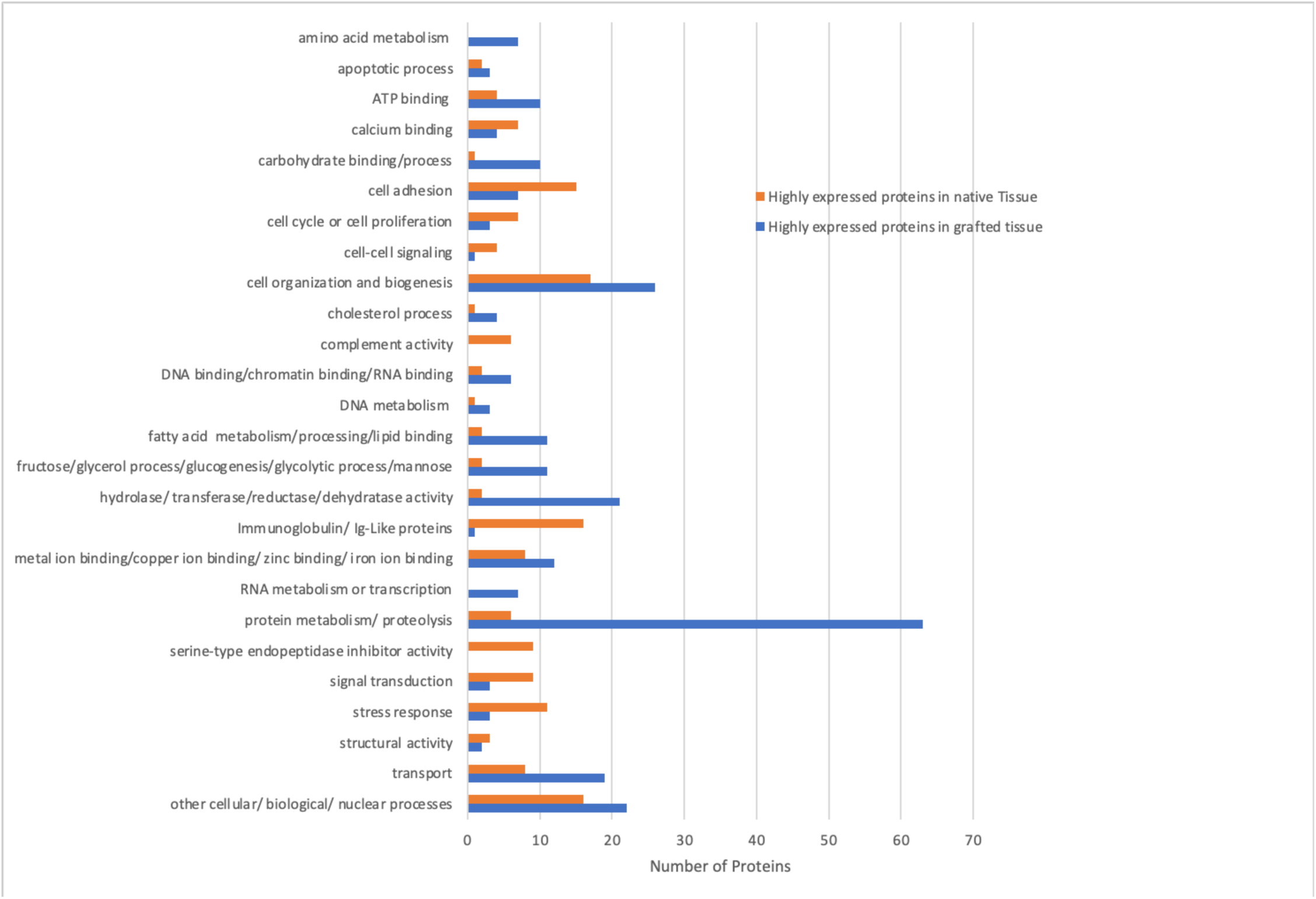
Biological Process Distribution for highly differentially expressed proteins in the enterocystoplasty study. The blue color-coded bars indicate the highly expressed proteins in the grafted tissue vs. the native tissue; while the orange coded bars indicate the highly expressed proteins in the native tissue vs. the grafted tissue.

## 4 DISCUSSION

In this study we have reported the proteomic profile of bladder augmented tissue in comparison to its native bladder tissue in a long-term study using a baboon bladder augmentation model. Three augmentation scenarios were employed that included the **E**-autograph ileal graft, the stem cell seeded polymeric biodegradable scaffold POCO, and the stem cell seeded biological scaffold SIS. The cells used for seeding onto the scaffold included autologous donor-matched bone marrow MSCs and CD34+ HSPCs. Over the course of the study in our baboon model, these cells would regenerate the portion of the cystectomized bladder. In our study, we used the cut off of log2 > 1 (or 2-fold with higher expression in the grafted or regenerated tissue) or log2 < −1 (or 2-fold with higher expression in the native tissue) and the significance level of p-value > 0.05 for the differentially expressed proteins in one tissue vs. the other. Using these criteria, we have found that the gold standard enterocystoplasty procedure that is clinically used yielded the most differentially expressed proteins in the grafted tissue vs the native tissue at a total of 419 proteins, with 259 proteins expressed at higher levels in the grafted tissue vs. native and 160 proteins expressed at higher levels in native vs. grafted tissue. For the proteins with higher expression in the native tissue vs. the grafted tissue included 4 proteins with log2 ratio of −3.13 to −4.09 (for a fold of 8.75 to 17.03), which included PCP4 (Purkinje cell protein 4), TRIM29 (Tripartite motif containing 29), AGTR2 (Type-2 angiotensin II receptor), and HPSE2 (Heparanase 2).

The proteins with the highest fold level of differential protein expression were in the ileal grafted tissue where a total of 65 proteins had a log2 ratio of 3.00 to 6.64 (or a fold of 8 to 99.73). The protein DEFA6, Defensin alpha 6, had the highest differential fold difference of log2 ratio of 6.64 or 99.73. DEFA6 is highly expressed in the secretory Paneth cells that reside in the small intestine [19,20]. DEFA6 protects the intestinal mucosa and defends against invasion of viruses and bacteria by forming fibrils and nanonets that encompasses pathogens [21]. It has however also reported that DEFA6 is also highly expressed in colorectal cancer (CRC) cell lines and patient samples [22]. By knocking down DEFA6 expression via shRNA in cancer cells, Jeong et al observed significantly inhibited cell growth, migration, and invasion in cancer cells in vitro, and inhibited tumorigenesis in vivo compared to control cells. Furthermore, it was determined that high DEFA6 expression to be a strong prognostic indicator for CRC as high expression of DEFA6 was observed in 51.4% (or 181/352) primary colorectal cancer tissue samples with high correlation to poor prognosis [21]. In a study by Husman et al, it was revealed that patients who undergone BAE had an incidence of 5-6% to develop bladder cancer after 50 years old [23]. It is worthy to note that whether or not that DEFA6 was observed to be highly expressed in cancer tissue that investigation into this protein maybe warranted to determine its potential role in tumorigenesis in bladder tissue post BAE.

In our study using the cell seeded graft, **CS-POCO** or **CS-SIS**, the proteomic analysis revealed the least number of differentially expressed proteins. **CS-SIS** had a total of 6 proteins including 1 protein having higher expression level in grafted vs. native at log2 ratio of 1.09 (2.13), and 5 proteins higher expression levels in native vs. grafted tissue and ranged from log2 ratio at −1.160 (fold of 2.23) to −1.270 (2.41). Our in-house **CS-POCO** graft yielded the least number of differentially expressed proteins with two proteins having higher expression in the native tissue vs. the grafted tissue. These two proteins were CHRDL1 (Chordin like 1) and LGALS2 (Galectin-2) which had log2 ratio at −1.030 (fold of 2.04) and −1.49 (fold of 2.81).

These data suggested that the ileal graft used in enterocystoplasty expressed many proteins that differed in the grafted tissue vs. native tissue, confirming a mismatch in tissue type. Using either **CS-POCO** or **CS-SIS** with the least of differentially expressed proteins in the regenerated tissue vs. native indicated better protein compatibility to the native tissue bladder. The data suggested further investigation into the use of CS-POCO or CS-SIS as potential cell seeded graft for the bladder augmentation as they support similar protein expression pattern compared to native bladder tissue.

## ACKNOWLEDGEMENTS

The authors would also like to acknowledge the Michelon Family and Legacy Healthcare (AKS) for their support and generosity.

